# Stochasticity in viral infection and host response: A competition between speed and reliability

**DOI:** 10.64898/2026.03.08.710362

**Authors:** Oskar Struer Lund, Ulrik Hvid, Bjarke Frost Nielsen, Kim Sneppen

**Affiliations:** Copenhagen University

## Abstract

The early stages of viral infection constitute a race between viral proliferation and interferon (IFN)-mediated defenses. Recent experiments on single-cell viral kinetics have demonstrated a high degree of stochasticity in the timing of viral release, but how this shapes the competition between virus and host remains unclear. We formulate a stochastic spatial model to address the question of how variability in the release of viral progeny and IFN affect the early infection dynamics. The model distinguishes between two types of timing noise: stochasticity in the initiation of release, and variability in the secretion time of individual virions. Our key result is an asymmetry in how noise affects outcomes: For the virus, stochastic initiation accelerates expansion, while for the host, effective containment via IFN benefits from precisely timed responses. For the secreting states, we find that a broader secretion profile (higher variability in particle release times) is always advantageous. In all cases, we find that stochasticity in signal timing plays a huge/central role in the early infections states.

## I. INTRODUCTION

Viruses are ubiquitous, infecting nearly all organisms and producing vast numbers of progeny per infected cell. Hosts counteract these dangerous predators; in mammals, early defenses include interferon (IFN) signaling and the activation of restriction factors [1]. Yet viruses persist through suppression of IFN signaling, exemplified by viruses as diverse as SARS-CoV-2 [2–4], Ebola [5] and Herpes [6]. The early stages of infection, before the activation of adaptive immunity, are heavily governed by a local tissue battle between the interferon and the virus. The fact that the battle is local is ignored in mean-field models [7]. Interestingly theese spatial effects results in a “ring-vaccination” where the IFN is able to confine the virus into a finite plaque. This is greatly discussed in [8], as well as the variability in IFN signalling. In this paper, we extensively focus on the effect of stochasticity in timing of viraland IFN signalling. Accordingly, a major part of the infection dynamics is local spreading, which can be modeled by stochastic cellular automata. Recent advances, such as single-cell sequencing [9–13] and spatial transcriptomics [14, 15] will, in the future, allow us to calibrate the modeled dynamics more carefully to observations of infection spreading in real tissues.

The fate of a single cell upon viral infection is governed by the ability of the virus to overcome intercellular defense mechanisms that otherwise repress viral entry and alarm neighboring cells via IFN secretion. Viruses have evolved numerous strategies to counteract cell defenses. Many RNA viruses inhibit the RIG-I pathway by blocking its adapter protein MAVS; for example, Influenza A virus uses its NS1 protein to interfere with the TRIM25-dependent ubiquitination of RIG-I, preventing full activation [17]. Rotaviruses encode NSP1, which indirectly inhibits transcription of interferon genes [18]. Similarly, coronaviruses such as SARS-CoV-2 antagonize multiple steps of the IFN pathway, including blockade of STAT1/2 signaling [19].

The diversity of viral evasion mechanisms illustrates that while cells may act as first responders, viruses in turn adapt by undermining this antiviral response. This process is highly stochastic, however, as only a minority of cells produce IFN in response to infection [20, 21], which is largely an all-or-nothing binary decision [20, 22].

This paper focuses on stochastic timing between different cell states and the outcome of infection. Cell-to-cell variability has multiple origins, including the local environment and the number of virus particles that infect a cell [23]. Here, we will concentrate on two aspects of variability: First, the time at which an infected cell starts to produce viruses/interferons, and secondly, how wide a temporal window the subsequent secretion happens over. The study is divided into two idealized scenarios.

First, we introduce the burst model, which focuses exclusively on stochastic fluctuations in the time to initiate viral or interferon production, by truncating all secretion to an instant event. In this formulation, once a cell reaches the particle-producing state, it immediately affects all neighboring cells within a prescribed radius *r*_*x*_. This burst model approximation isolates the impact of early-stage timing variability upon secretion, while capturing the instant release of the virus for cytolytic (“lytic”) viruses, including poliovirus [24], adenovirus infection in epithelial cells [25–27], and many non-enveloped viruses [28].

Second, we consider a shedding scenario in which production begins after a fixed initial delay, allowing us to isolate the role of the secretion-phase duration. Here the temporal profile of secretion modulates infection dynamics by shaping both the effective burst size and the spatiotemporal distribution of released particles. In contrast to the instantaneous-burst approximation, we explicitly prescribe a burst size together with a space–time secretion profile. For clarity, we suppress stochasticity in the earlier intracellular stages leading up to the release.

These two models can be thought of as representing two different biological scenarios (lytic vs. gradual virion release) or as two different idealizations of the same underlying process, with the burst model emphasizing cell-to-cell variation in viral generation time, and the shedding scenario addressing the significance of the viral release occurring over time.

Throughout the paper, we will employ Gamma distributions for both delay times (burst model) and secretion profiles (shedding scenario). Fig. 1a illustrates the time distributions used to fit both intracellular transitions and secretion density profiles:

**FIG. 1.**
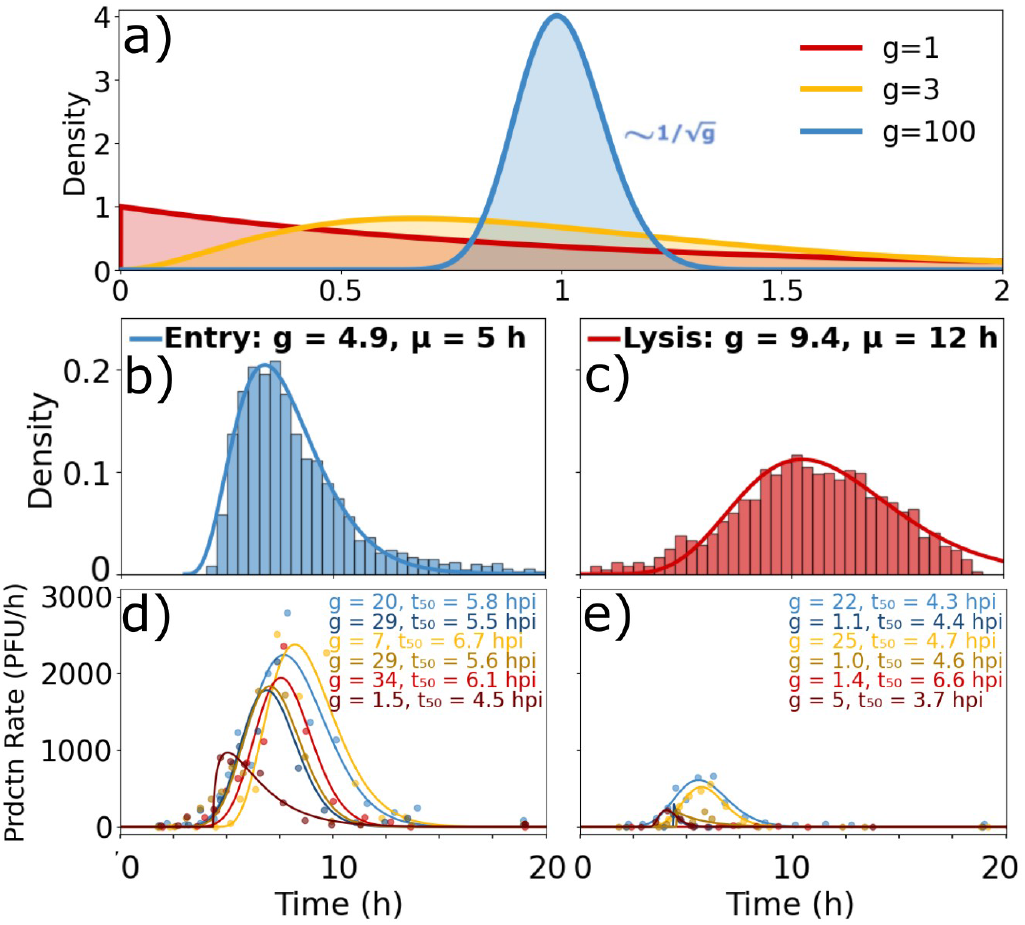
Kinetics of viral release. **a)** Gamma distributions with the same average but different shape parameters, *g*. The distribution is used to fit viral release kinetics in this study. **b)** Probability density of entry (infection) time for lytic poliovirus, using data from [9]. **c)** Probability density of lysis time of poliovirus counted after viral entry. Importantly, ~50% of the cells never lysed, hence this distribution shows the lysis times of those cells that actually lysed within the time frame of 24 hours. Entryand lysis times are inferred from viral mRNA concentration data in cells infected by a Polio virus, sampled across ~3000 cells. The blue histogram shows the time needed for a virus to infect an epithelial cell in a trapping region ~3 times larger than the cell. The cell is exposed to poliovirus for 1 hour under conditions giving an average MOI of ~5. The red histogram shows the lysis time, counted from viral entry. **d**,**e)** Observed single-cell virus production rates for Vesicular stomatitis virus (VSV) as a function of time since introduction to a culture of BHK cells [16]. The continuous lines illustrate Gamma distribution fits obtained by non-linear least squares (see supplement). *t*_50_ denotes the time until half of the virus are released. This virus does not lyse the cell but secretes progeny over a prolonged period. See supplement (Fig.S8) for raw single-cell data.

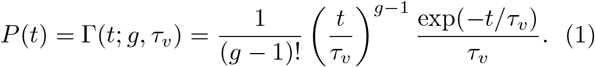

In this Gamma distribution, a small shape parameter *g* reflects a wide distribution of waiting times, while a larger *g* implies more deterministic behavior, with a standard deviation 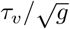.

## II. DATA AND MODEL CONSIDERATIONS

Using data from [9] Fig. 1b–c) illustrates a pronounced cell-to-cell variability for a lytic virus, in which all progeny virions are released upon host-cell rupture. Figure 1c shows the distribution of times from viral entry to cell lysis (the latency time). These latency times are well described by a Gamma distribution with shape parameter *g* ≈10, corresponding to a coefficient of variation of approximately 30%.

Figs. 1d-e show examples of time courses of virus yield for a virus (Vesicular stomatitis virus, VSV) that does not burst its mammalian host cell. Instead, it induces a prolonged emission of virus particles from the cell, starting about 5 hours after infection and gradually ending after about 15 hours. The total number of virions released varies substantially between cells, with a mean of ≈ 4000 virions, ranging from approx 100 to 10^4^ virions. The Gamma distribution differs substantially from cell to cell, with the form factor *g* taking values between 2 to 34, and a peak rate between approx. 200 and 2500. A typical cell releases viruses for around 10h with *g* ≈10.

Figure 1 reveals substantial heterogeneity in both infection timing and total virus production; however, how this variability shapes infection spread within mammalian tissue remains unresolved. Before introducing the full model, we first illustrate, in a simplified setting, why it is essential to incorporate realistic levels of stochasticity. For simplicity, we consider the burst model. First we consider a virus spreading through naive tissue that does not mount an interferon response. This would represent infection of Baby Hamster Kidney (BHK) cells. The tissue is implemented as a two-dimensional square lattice, representing an epithelial sheet (Fig. 2). After a cell is infected, the virus must replicate intracellularly before progeny virions are released. We assume this replication requires, on average, a latency time *τ*_*v*_, which in Fig. 1d,e is approximately six hours [16]. In the burst model, this viral release is assigned to a single burst that immediately infects all susceptible cells within a radius *r*_*v*_.

**FIG. 2.**
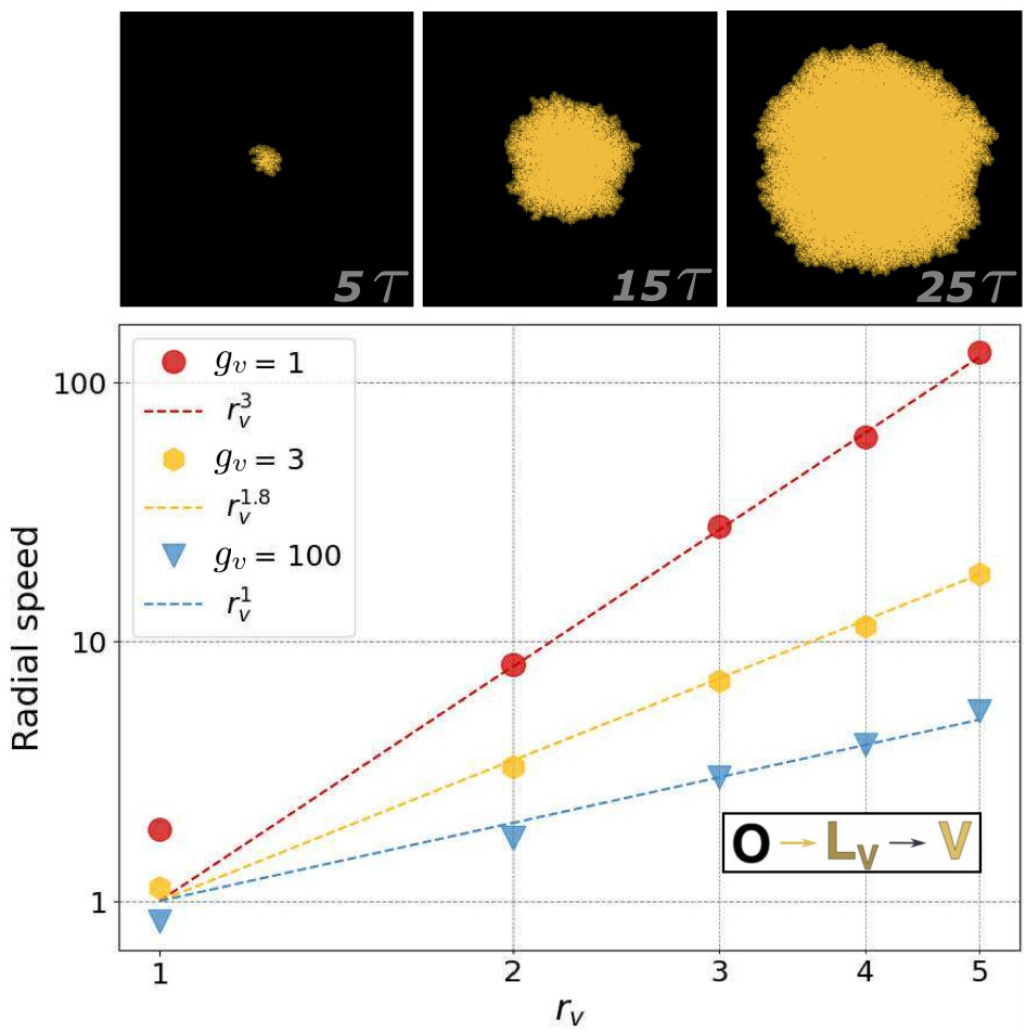
Stochastic transitions favor fast spreading. The top row illustrates a highly stochastic virus, 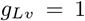, that spreads with *r*_*v*_ = 3 from 5*τ* till 25*τ* on a lattice of size *L* = 300. The lower log-log plot shows the speed of plaque expansion as a function of *r*_*v*_ for different values of 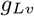. When 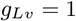 (red), rare fast transitions dominate and speed scales approximately as 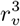. For 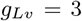 (yellow), expansion slows (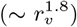). In a more deterministic context (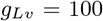, blue), speeds scale roughly linearly with *r*_*v*_.

Figure 2 depicts viral spread in this interferon-deficient tissue model. The lower panel shows that the dependence of the invasion speed *v* on the infection radius *r*_*v*_ is highly sensitive to heterogeneity in burst (lysis) times. Specifically,

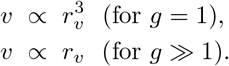

The mechanistic origin of this contrast is a consequence of the wide waiting time profiles upon bursting. For exponentially distributed waiting times (*g* = 1), the advance of the infection front is governed by the *earliest* burst among the 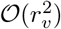 newly infected neighbors. The expected minimum latency, therefore, decreases with neighborhood size, 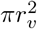 scaling as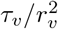. Because the characteristic spatial advance during this time is ~ *r*_*v*_, the resulting front propagation speed scales as 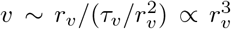 with *τ*_*v*_ = 1 used throughout this paper. In the opposite limit of large *g*, burst times are narrowly distributed and effectively deterministic, thus the propagation speed will be *r*_*v*_*/τ*_*v*_ = *r*_*v*_.

## III. MODELS AND RESULTS

We represent the epithelial tissue as a cellular lattice with different procrastination routes to three final states: Viral, IFN-producing, and antiviral state. This is illustrated in Fig. 3. The final fate of each cell is governed by the timing and type of received signals. Following viral entry, the cell progresses to a viral-producing state, unless it receives an IFN signal. Conversely, exposure to IFN will make the cell enter a dormant antiviral state [29]. However, if the cell on the way to the antiviral state becomes infected, then it instead becomes IFNproducing. Hence, IFN (*N*) and virus (*V*) producing cells are competing for local tissue dominance. All states and transitions are summarized in Fig. 3, and the associated waiting times are modeled using Gamma-distributed delays.

**FIG. 3.**
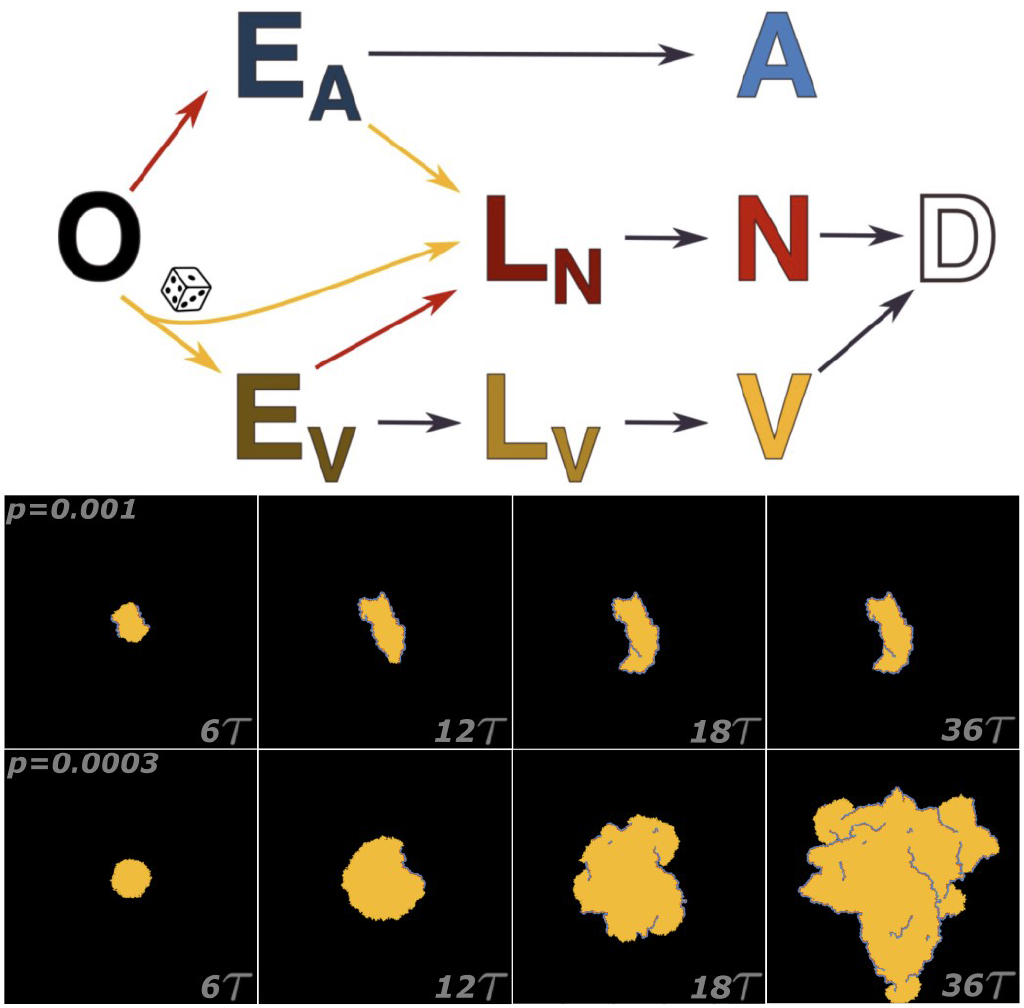
Model: Naive cells (*O*) exclusively exposed to the virus follow the sequence 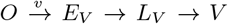, while IFN exposure triggers *O E*_*A*_ *A* [31] and co-occurring, within time frame *τ~* 6 hours, virus and IFN favors *O → E*_*N*_ *→ N* [32, 33]. Yellow arrows represent transitions driven by virions, while red arrows denote IFN-driven transitions. Both occur instantaneously upon receiving their respective signals. Black arrows indicate intracellular processes that proceed stochastically over time. This stochasticity is modeled by Erlang-distributed delays, parameterized by *g*_*v*_ and *g*_*n*_, which control the variability of viral- and IFN-induced transitions, respectively. The mean transition time between states *E*_*V*_ *→ L*_*V*_ and *E*_*A*_ *→ A* is *τ*. Importantly, there is a fundamental difference between *E*-states and *L*-states: In *L*-states, the cell is irreversibly committed to its final state. In *E*-states, the final faith of the cell can still be influences of either virus-or IFN signals. These *E*- and *L*- states are motivated from [23]. The *L*_*N*_ state has a mean duration of 2*τ*, making the mean time to reach *N* and *V* state the same. The dice symbol indicates this small probability *p* of a cell being a first responder in the absence of IFN stimulation. In this model, *E*_*V*_ and *L*_*V*_ share the same stochasticity parameter *g*_*v*_, while *E*_*A*_ and *L*_*N*_ share *g*_*n*_. Lower panels: Snapshots of two simulations with identical parameters (*r*_*v*_ = 1, *r*_*n*_ = 3, *g*_*v*_ = 1, *g*_*n*_ = 100) but different fractions of first responders (*p* values). In the upper row, *p* = 10^−3^ leads to confinement, whereas *p* = 3 · 10^−4^ in the lower panel leads to unlimited spreading.

Cells exposed to the virus will activate an IFN response with probability *p*, which in our model parameterizes the stochastic nature of activation of first responders. This mimics a process that is largely driven by cytosolic sensors such as RIG-I, which detect viral RNA and signal the cell to produce IFN [30]. Hence, the first IFN response will appear when the infection size is on order ~1*/p*. Depending on the fraction of first responders (*p*), the range of IFN signals (*r*_*n*_), the range of viral diffusion (*r*_*v*_), and the shape factor of the Gamma distribution, we observe either expansion or containment of infection.

### A. Burst Model

In this formulation, both infection and signaling are modeled via a local radius-of-influence rule. At each time step, virus-producing (*V*) cells and interferon-emitting (*N*) cells act on all lattice sites within fixed interaction radii: *r*_*v*_ for viral transmission and *r*_*n*_ for IFN signaling. Concretely, around each emitting cell, we define a disk of radius *r*_*v*_ or *r*_*n*_ and label all sites within the disk as exposed to the associated signal, virus or IFN. The lifetime *τ* of exposed states would be of order ~6 hours, see [23].

The radius-of-influence formulation provides a coarse-grained representation of a lysis-like process: Instead of tracking the diffusion of individual virions or IFN molecules, each active cell instantaneously communicates with its neighborhood within the specified range. This is simplification is most justified for the small interferon molecules, but less for a larger virus in the limited space between cells. Our simple, geometric rule emphasizes the competition between viral expansion and IFN-mediated containment. The burst model focuses on the importance of the time distribution to reach a viral or an interferon-producing state measured in containment probability, rather than focusing on the underlying secretion process. We address the influence of the secretion process itself in the “Shedding scenario” section.

The speed of the interferon response is also governed by stochasticity; however, with the constraint that IFN can only activate itself in the narrow band of *E*_*V*_ -tissue at the edge of the infection plaque. The characteristic width of this frontier is set by the distance traveled by the infection front during the eclipse time, 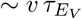. Therefore, variability yields at most a modest enhancement of the effective IFN spread speed, see Fig. S6.

Primarily, containment of infection hinges on maintaining the integrity of the IFN barrier. As illustrated in Fig. 4a,b, this barrier is only a few cells thick, making it inherently vulnerable to fluctuations. With stochastic activation of IFN 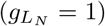, the resulting variability generates local gaps in this thin layer, through which the virus can escape. The associated fragility requieres a significant higher fraction of first responders to ensure confinement: Reliable containment requires *p* ≳ 10^−2^ under very noisy 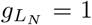 IFN activation, whereas a more temporally precise IFN response can achieve containment with far fewer first responders, on the order of *p* ≳ 10^−4^, see 4d).

**FIG. 4.**
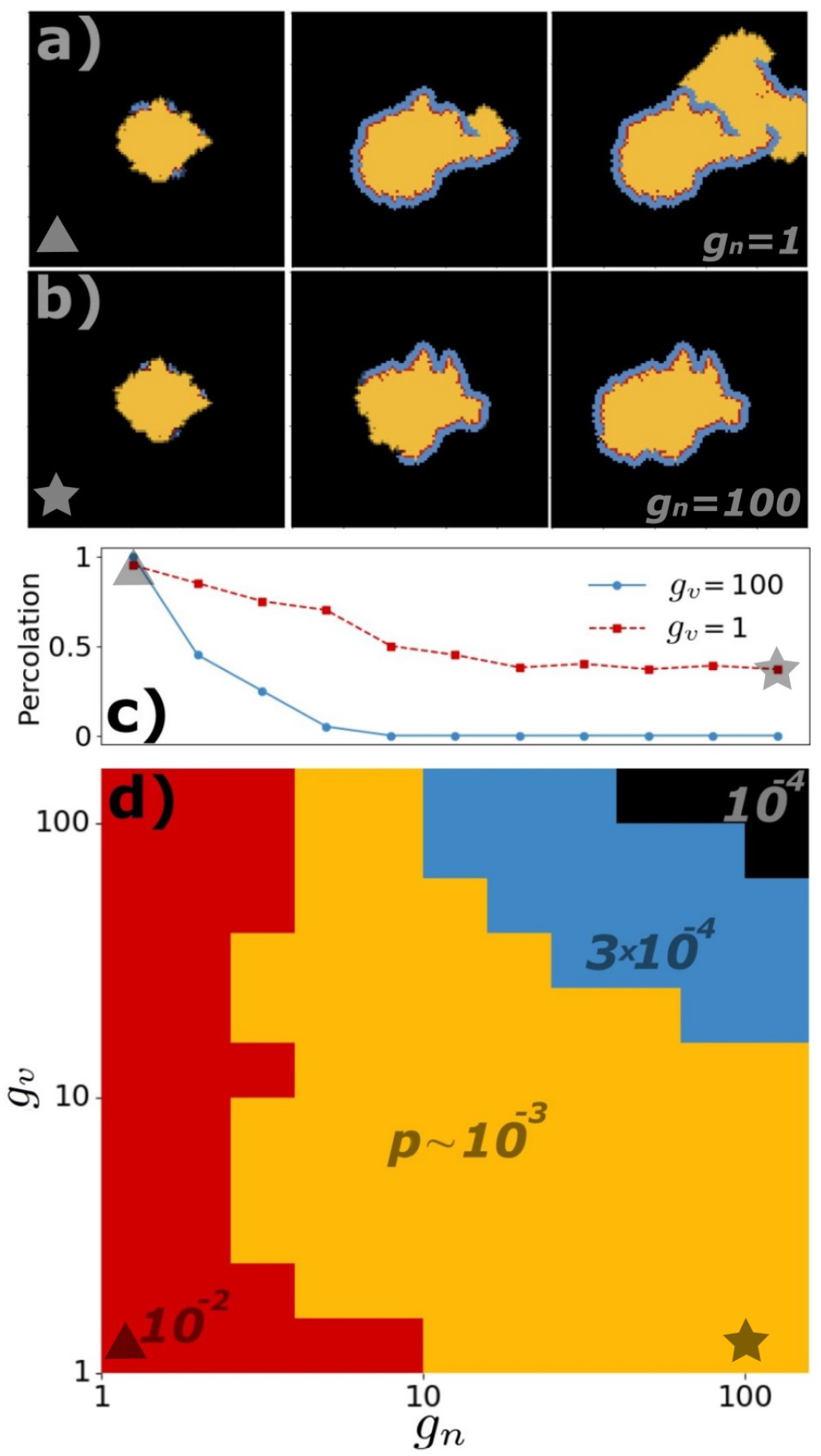
Viral stochasticity and IFN precision in the burst model. Across all simulations, we set *r*_*v*_ = 1 and *r*_*n*_ = 3. **(a**,**b)** Snapshots with *p* = 0.003. Both panels use stochastic viral dynamics (*g*_*v*_ = 1 for both *E*_*V*_ and *L*_*V*_ states). In (a), stochastic IFN (*g*_*n*_ = 1) fails to contain viral spread, whereas in (b), a more deterministic IFN (*g*_*n*_ = 100) succeeds in confining the virus, illustrating the importance of reliable IFN signaling. If IFN-signalling is noisy, random gaps allow viral escape. **(c)** Percolation probability on a *L* = 1000 lattice over 1000 runs with first responder fraction, *p* = 10^−3^. Percolation is defined by the viral plaque reaching the lattice edge. Stochastic virus (*g*_*v*_ = 1, red) percolates more often than deterministic virus (*g*_*v*_ = 100, blue). **(d)** First-responder fraction *p* among values *p* = 10^−4^, 3 · 10^−3^, 10^−3^, 10^−2^ required for 50% or more containment as a function of *g*_*n*_ and *g*_*v*_ (lattice size *L* = 1000). Viral stochasticity (*g*_*v*_ = 1) promotes spread, while IFN precision (*g*_*n*_ = 100) enhances containment.

Thus, a fundamental asymmetry emerges: the virus benefits from stochasticity in the time required to reach the virus-producing state accelerating, its spatial spread.

In contrast, the host benefits from a precise reliable response, preserving the integrity of IFN-wall. This asymmetry is quantified in Fig. 4: for small *g*_*v*_, a larger *p* is required to contain an outbreak, while for larger *g*_*n*_, a smaller *p* is sufficient.

### B. Shedding scenario (duration of secretion)

We now relax the assumption that *V* - and *N* -signals are released in instantaneous bursts, and instead model actual secretion over time mimicking the shedding process from infected cells seen for viruses such as SARS-CoV-2, influenza [34, 35]. This allows us to examine the consequences of the early release of a small fraction of virions or interferon molecules.

Concretely, we assign virus-producing and interferon-producing cells the parameters (*g*^*Lx*^, *Y*_*x*_, ⟨ *r*_*x*_ ⟩), where *x* = *V or N*, and:

- *g*_*Lx*_ determines the shape of the temporal release profile from the latency state *L*_*>x*_,
- *Y*_*x*_ is the total yield,
- ⟨ *r*_*x*_ ⟩ is the mean particle displacement before absorption by a cell.

For a cell in state *L*_*x*_, the number of particles released during a time interval [*t, t* + d*t*] is:

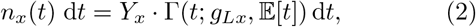

where Γ(*t*; *g*_*Lx*_,𝔼 [*t*]) is a Gamma distribution of *t* with parameters *g*_*Lx*_ (shape) and 𝔼 [*t*] (mean). *t* = 0 corresponds to the transition into the secreting state, and *Y*_*x*_ is the total yield of signal particles sent, which for interferons is counted in units of the minimal amount needed to trigger a response in a receiving cell.

Each of the *n*_*x*_(*t*) d*t* particles is subsequently placed in a random direction at a distance *r* according to the distribution:

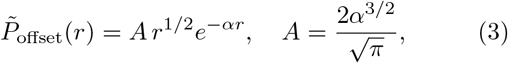

such that the probability for a virion to land at a distance between *r* and *r* + d*r* is 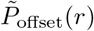 d*r*. The parameter *α* is related to the mean displacement via ⟨*r*⟩ = 3*/*(2*α*).

For simplicity, we let 𝔼 [*t*_*n*_] = 𝔼 [*t*_*v*_] = *τ*, and set the mean durations of the exposed states *E*_*V*_ and *E*_*N*_ to *τ* as well, with high precision (*g*_*E*_ = 100), prior to progressing into *L*_*V*_ or *L*_*N*_. We thus keep the time to enter the particle-producing state fixed, allowing us to focus on the effects of varying the secretion profile shape. The nearly deterministic initial delays also prevent a fraction of cells from transitioning almost immediately to viral or interferon secretion, which would otherwise lead to extremely rapid propagation, as explored in the burst model.

The shedding scenario thus highlights a new aspect of the trade-off between precision and speed, by allowing us to examine the distinction between precision in the secretion profile, as opposed to the *initiation* of signaling that was addressed via the burst model. Fig. 5b,c illustrates the superiority of a fast, rather than precise, response. Panels d and e show that the number of cells hit over time for a single particle-secreting cell, with the red curve describing the fast and noisy response, while the blue curve shows the slower but precise response. Panels b and c clearly indicate that the red, noisy secretion profile is superior for the IFN as well.

**FIG. 5.**
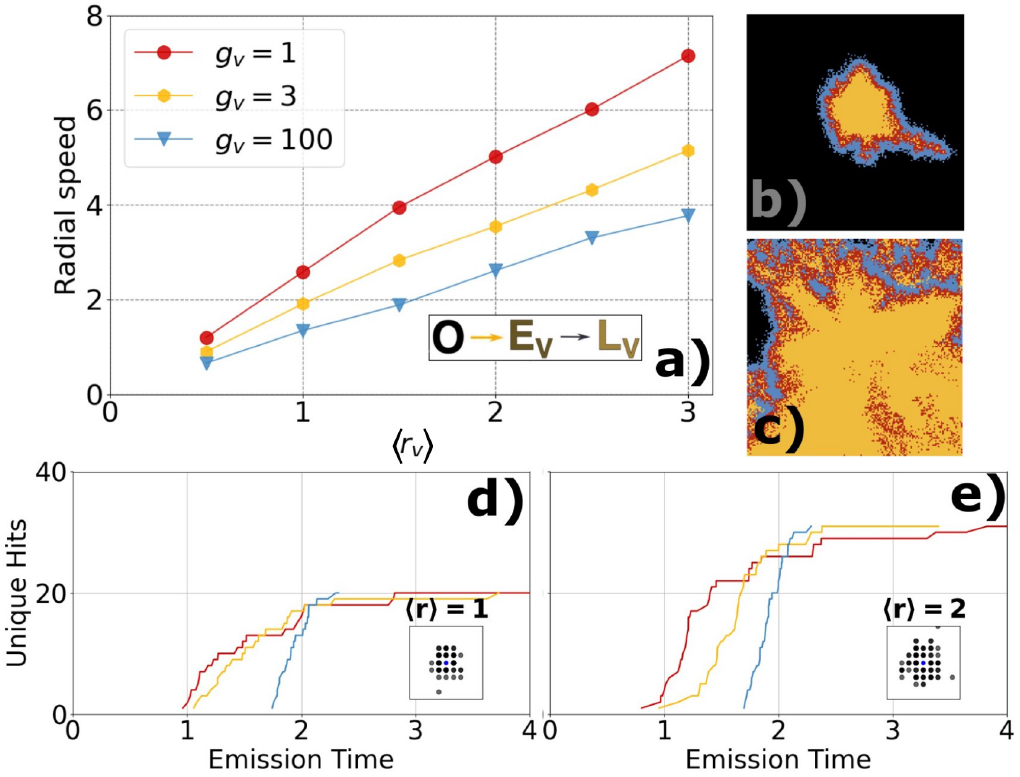
Trade-off: Speed vs. precision in Shedding scenario. Here we fix the number of Erlang gates in EV at 100 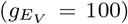 and insert an intermediate state E_N_ with 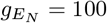, before the cell progresses to L_*N*_. This suppresses stochasticity in the time upon secreting states are reached. We vary the number of Erlang gates in the secreting states, L_V_, L_N_ via shape parameters, g_*v*_, g_*n*_. The burst size is fixed at 100 for both interferon and virus, while fig. S1 examine burstsize N_*x*_ = 10^3^. a) Plaque expansion speed as a function of r_v_ describing the mean square distance of virus infections. Stochastic viral secretion (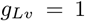, red) leads to faster spread than more deterministic secretion (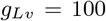, blue), since some cells begin producing virus much earlier. This is further examined in Fig. S3, which quantifies the time required to infect 50%. b,c) Explore infection outcomes for fixed p = 0.001, ⟨r_v_⟩ = 1,⟨, r_*n*_⟩ = 2. b, interferon secretion is broad and stochastic 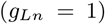, while panel c use 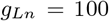. The comparison highlights the consequences of slow but precise IFN responses, which allow viral spread to escape before the antiviral barrier is established. d, e) Number of unique neighbor cells reached by a burst of 100 particles, with mean squared displacement ⟨r⟩ = 1, and ⟨r⟩ = 2, respectively. Fig. S3, shows how unique neighbors scale with burst size. These results apply to both viral- and interferon particles.

Fig. S3 illustrates how the total number of hits scales with the particle yield and compares the spatiotemporal influence of noisy and precise secretion profiles. The substantial cell-to-cell variability in *g* seen in the experiments of ref. [16] (see Fig. 1) raise the possibility that a small subpopulation of “fast” cells disproportionately accelerates spatial spread. Such front-driven fluctuations may be further amplified if induced antiviral states impose heterogeneous infection thresholds across the tissue.

The shedding scenario predicts that temporally precise IFN release (a narrow IFN profile) is a substantial *dis*advantage. The stark contrast between Fig. 5b (where 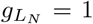) and Fig. 5c (where 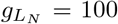) shows that tightly timed IFN secretion fails to halt viral spread, whereas gradual production, which includes an early IFN component, succeeds. This behavior is quantified in Fig. 5d,e: the noisy (red) trajectories reach a large fraction of the accessible neighborhood rapidly, highlighting the importance of early signaling in establishing protection ahead of the advancing infection front. However, the interferon still needs to be precise/reliable in *entering* the secreting state, see Fig. S5, agreeing with the main insight of the burst model.

For both Virus and IFN, the stochastic release rapidly reaches the majority of its total targets. Fast responses are especially effective because neighbors who are activated early can themselves begin signaling sooner; thus, reaching half of the neighbors quickly is arguably more important than reaching distant neighbors.

In the Shedding scenario, we don’t observe any dramatic superlinear scaling of expansion speed with the viral release radius seen in the burst model (compare Fig. 5a with Fig. 2b), that resulted from the scaling of the viral load with 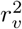 inherent to the burst model. The absence of a superlinear scaling is due to two reasons. In the burst model, we increased the range of the area hit. That means both mean square distance displacement *and* viral yield would be needed to have a similar effect in the shedding scenario. In this setup, we only increase the mean square virus displacement. Further, we eliminate the chance of extremely early virus-producing cells by introducing the nearly deterministic *E*_*V*_ -state.

## IV. DISCUSSION

Our study combines two complementary scenarios for the interplay between viral spread and IFN signaling in epithelial tissue. The burst model offers a minimal description of a lytic virus, where infection and IFN-signaling are represented as circles. This framework explores the critical role of stochasticity in the *timing of transitioning to a virus/interferon producing state*, illustrating the asymmetry between optimal viral- and interferon behavior; viruses benefit from stochasticity, and interferon benefits from precision in the time to progress to the emitting state.

The shedding scenario introduces an additional parameter, the total yield, with a particle-based formulation of both virus and interferon propagation. In this formulation, fixing the time to reach secreting states, the optimal strategy for the virus and its host is shared. Fast signaling becomes optimal, with the key intuition that the need to quickly alert close neighbors outweighs the need to alarm all neighbors at some later time. Meanwhile, the close neighbors can begin to alarm *their* neighbors. The shedding scenario predicts that the optimal secretion strategy is fast and incomplete emission for both virus and interferon.

Together, these two scenarios show that infection outcomes at the tissue level are shaped by both stochasticity and coordination. The interplay between viral noise and the precision of innate IFN responses remains relatively underexplored experimentally, and our findings may contribute to facilitating further research into the dynamics governing local infection dynamics. Future work could extend this framework by explicitly incorporating variability in burst size (Fig. 1b,c) and by determining Lévy flight-like characteristics of viral movement through epithelial tissue, driven, for example, by occasionally accessing small blood vessels. As noted in the introduction, viral expansion during the early stages of infection in a host is much slower than the burst size of individual cells would naively imply, implying that successful long-range dispersal events are rare, occurring on the order of one such jump per infected cell.

Other aspects that were ignored in our model include indirect cooperativity between virions [36]. This could be addressed in future extensions of the shedding model by allowing higher-MOI infections to exhibit different burst sizes. In addition, infections with defective virus particles may add further complexity to the virus-producing and interferon-producing branches of Fig. 3a [37].

In summary, our simulations support three conclusions: First, viral expansion is promoted by *stochasticity* in the time to reach the virus-producing state: noisy intracellular kinetics accelerate plaque growth. Second, effective immune containment exhibits the opposite requirement: robust control relies on *temporal precision* in the transition to interferon production, because reliability is essential to maintain an intact protective “containment ring”. Third, despite this asymmetry, both the IFN and the virus benefit from rapid start of secretion: fast, noisy, and partial signaling can outperform slow, precisely timed, massive but persistently delayed release. Overall, these results emphasize that the outcome is often decided by which side first gains access to the advancing frontier and thereby wins the local race to establish infection or protection.

## Acknowledgment

This research was funded by the Danish National Research Foundation (grant no. DNRF170).

