## Supplementary Figures and Tables for "Stochasticity in viral infection and host response: A competition between speed and reliability"

### SUPPLEMENTARY INFORMATION

#### 1. Introduction

### SUPPLEMENTARY INFORMATION

This supplementary material provides extended analyses and data to support the findings in the main text. Section S1 details the statistical methods used to fit Gamma distributions to the single-cell viral kinetics data presented in Fig. 1. Section S2 presents additional simulations for the *shedding scenario*. Figure S2 demonstrates the necessity of a precise pre-exposed state by comparing radial spread speeds with and without a regulated delay. Figure S3 explores the fundamental trade-offs in signal coverage and timing as a function of burst size and secretion stochasticity. Figure S1M replicates the analysis of Fig. 5 from the main text but with an increased burst size of $10^3$ for both virus and interferon. Figure S5 underscores the critical importance of temporal precision in entering the interferon-producing state for establishing a reliable antiviral barrier. Section S3 contains an analysis for the *burst model*. Figure S6 illustrates how the effective propagation speed of the interferon signal depends on the spatial scale of viral spread ($r_v$), reaching its maximum only when $r_v \sim r_n$. Finally, Section S4 presents the raw single-cell data underlying the kinetic profiles in the main text: Figure S7 shows the dynamic intracellular viral RNA time series for lytic poliovirus, and Figure S8 displays the production rates for non-lytic VSV.

#### Fitting for fig 1

Let the observed data consist of time points $x_i \in \mathbb{R}_+$ and measured production rates $y_i \in \mathbb{R}_+$ for $i = 1, \ldots, N$. We assume the underlying kinetics can be represented by a shifted and scaled Gamma distribution of the form

$$f(x; k, \lambda, A, x_0) \; = \; \begin{cases} A \, \dfrac{\lambda^k}{\Gamma(k)} \, (x - x_0)^{k-1} e^{-\lambda(x - x_0)}, & x > x_0, \\ 0, & x \leq x_0, \end{cases}$$

where

- $k > 1$ is the *shape parameter*,
- $\lambda > 0$ is the *rate parameter*,
- $A > 0$ is an *amplitude scaling factor*,
- $x_0 \geq 0$ is a *time shift* (onset of production),
- $\Gamma(\cdot)$ denotes the Gamma distribution.

For each dataset, the parameters $\theta = (k, \lambda, A, x_0)$ were estimated by solving the nonlinear least-squares problem

$$\hat{\theta} \; = \; \arg\min_{\theta \in \Omega} \sum_{i=1}^{N} \left(y_i - f(x_i; \theta)\right)^2 ,$$

subject to the parameter bounds

$$\Omega = \Big\{ (k, \lambda, A, x_0) \;\Big|\; 1 \leq k \leq 10^2, \; 0 \leq \lambda \leq 5, \; A \geq 0, \; 0 \leq x_0 \leq 5 \Big\}.$$

The optimization was carried out using a trust-region reflective algorithm for nonlinear least squares (implemented in `scipy.optimize.curve_fit`). Initial guesses for $(k, x_0)$ were chosen based on expected rise times and skewness of the data, while $A$ was initialized from $\max_i y_i$ and $\lambda$ set to 1.

The fitted function $\hat{f}(x) = f(x; \hat{\theta})$ was plotted alongside the observed data points $(x_i, y_i)$. For visualization, the estimated shape parameter $\hat{k}$ was reported in each subplot as an indicator of the temporal sharpness of the response.

#### 2. Additional "*Shedding-Scenario*" Figures

The additional "*Shedding-Scenario*" figures are included to illustrate the need for a frozen exposed state, $E_V, E_N$ with a precise profile, $g_E = 100$, see Fig S2. Further this section also illustrates the characteristics of a precise profile and a noisy profile in terms of influencing neighbors, see Fig S3. This section also demonstrates the results robustness of main fig 5, repeated in Fig S1m with burst sizes of virus and interferon on $10^3$.
Figure fig. S5 illustrates that the need for precision in pre-particle producing state for the interferon is needed, to put up a reliable defense.

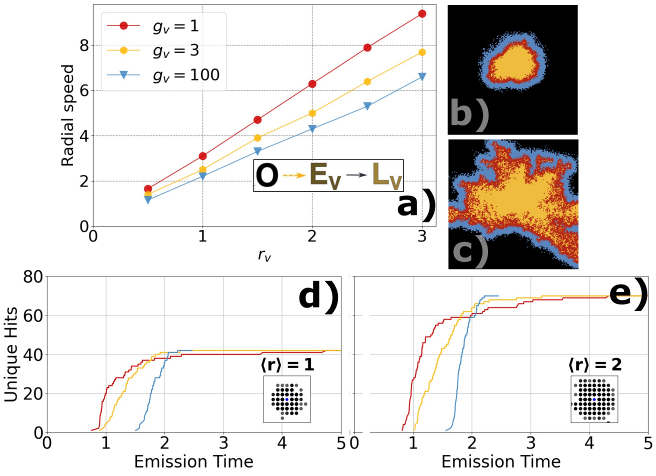


FIG. S1. **Speed vs. precision trade-off.** Burst size = $10^3$ across all simulations for both IFN and virus is fixed. **a)** Plaque expansion speed vs. $r_v$ for different Erlang gates. More random viral timing (1 gate, red) spreads faster than more deterministic timing (100 gates, blue) because some cells start producing virus much earlier. Right-side images compare IFN strategies: top, random IFN (1 gate); bottom, precise IFN (100 gates). Here, random IFN acts sooner and works better. **b–c)** When and how far particles spread: random IFN reaches more cells earlier; precise IFN spreads its effect more slowly. Panel **c** uses a larger spread radius than **b**.

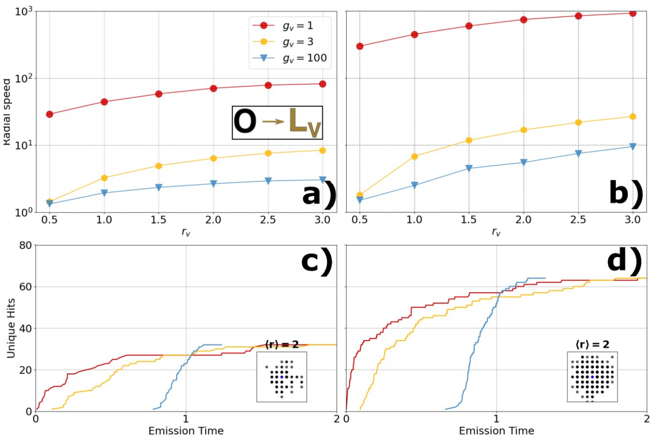


FIG. S2. **(a)** and **(b)** show radial speed versus $r_v$ for gate counts $k = \{1, 3, 100\}$ (red, green, blue). Panel (a) uses burst size 100; panel (b) uses 1000. **(c)** and **(d)** show the corresponding particle-release trajectories when the EV state is removed (immediate release). Across conditions, radial speed increases roughly linearly with burst size and $r_v$: the red ($k = 1$) curve scales at about, $30\% \cdot \beta_V \cdot r_v$, the green ($k = 3$) at about $2\% \cdot \beta_V \cdot r_v$, and the blue ($k = 100$) at about $1\% \cdot \beta_V \cdot r_v$, with $\beta_v := ViralBurstSize$.

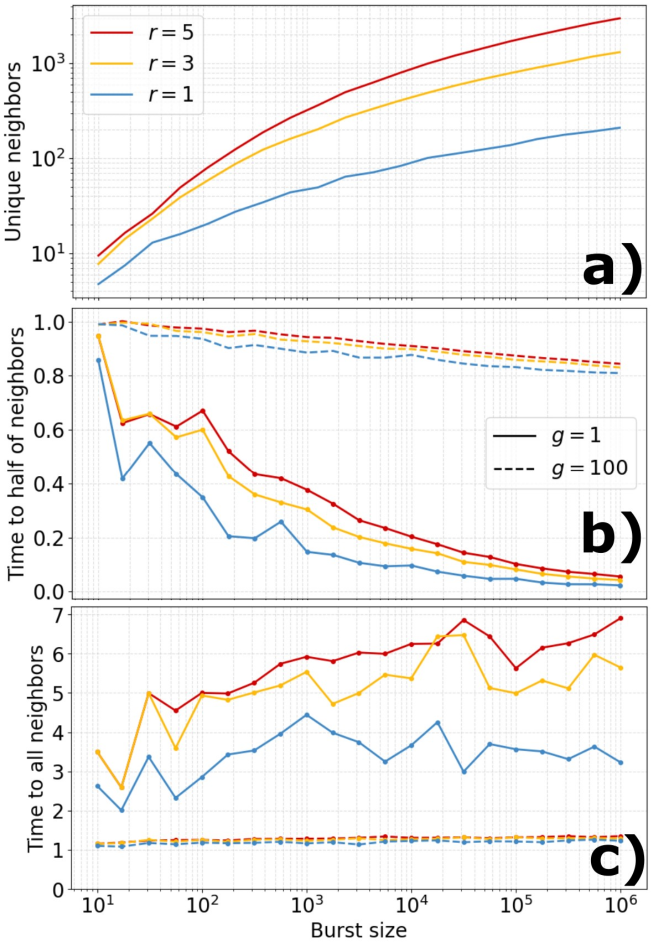


FIG. S3. This figure examines the relationship between secretory burst characteristics and the time required to alert neighboring cells. Panel (a) shows that for a fixed mean square diffusion distance, the number of cells contacted increases with burst size. Panel (b) compares the time required to reach half of all neighboring cells for different secretion Gamma distribution shape parameters. The dashed line represents a precise secretion profile ($g = 100$), which requires approximately $0.9\tau$ to contact half of its neighbors. In contrast, a noisy secretion profile ($g = 1$) achieves the same coverage in significantly less time; for a burst size $Y \sim 10^3$, the required time is roughly half that of the precise shedding scenario. Panel (c) examines the time needed to contact *all* neighboring cells. Here, the precise secretion profile ($g = 100$) outperforms the noisy profile ($g = 1$), reaching all neighbors more quickly. We observe that both the virus and the interferon benefits from having a noisy secretion profile, in conclussion, hitting neighbors fast and letting them reproduce signal outweigh the importance of hitting all neighbors.

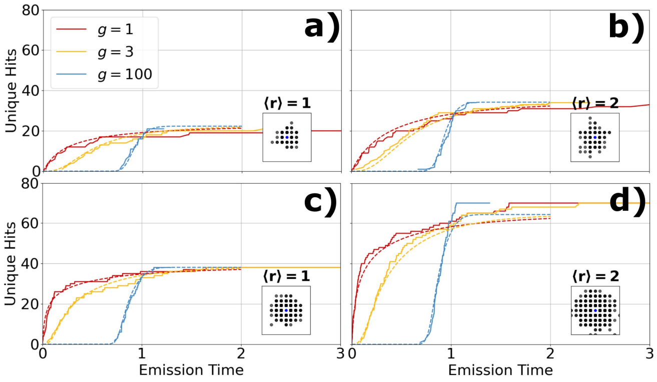


FIG. S4. **Analytical Fits.** Dashed colored lines follow the analytical curves for different gate realizations, $g = 1, 3, 100$, of particle releases over time. **a),b)** show results with different $\langle r \rangle = 1, 2$ and burst size $10^2$. **c),d)** are similar, but with burst size $10^3$.

For simplicity, there is no release delay here. Normally, we simulate with a pre-state of 100 gates upon releasing in latent states. Adding such a state would correspond to shifting all curves by $\sim 1$ to the right on the x-axis, simulating a delay of $\sim 1\tau$.

The dashed analytical curves show the expected number of unique sites visited. For each shape parameter $g = k$, the emission times follow the Gamma distribution with cumulative probability $q(t)$. The chance that a given site has not been visited by time $t$ is $(1-q(t)p_{ij})^N$, where $p_{ij}$ is the single-particle landing probability and $N$ the number of particles. Summing over all sites gives the expected number of distinct hits, $\sum_{i,j}\left[1-\left(1-q(t)p_{ij}\right)^N\right]$.

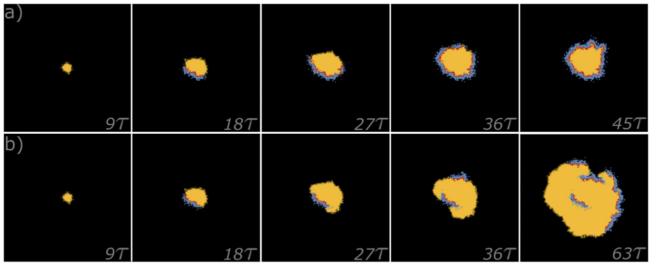


FIG. S5. **Temporal precision in interferon secretion timing determines the outcome of viral spread.** Both simulations use identical parameters: random seed; first-responder probability $p = 0.003$; mean emission distances $\langle r_v \rangle = 0.5$, $\langle r_n \rangle = 1.5$; burst sizes $\beta_v = \beta_n = 100$. Time spent in secreting states is tightly regulated, with $g_{Lv} = g_{Ln} = 100$, corresponding to a mean secretion duration of $1\tau$ in both states. Panel (a) shows that precise EN transition timing ($g_{EN} \sim 100$) leads to confined infection spread. Panel (b) shows that noisy EN transition timing ($g_{EN} \sim 3$) allows the infection to percolate across the grid. These results demonstrate that reliable, temporally precise transition into the interferon-producing state is essential for effective viral containment. This indicates that the principles derived from the burst model regarding precise $E_N$ timing also apply to the burst model, confirming the coherence of the findings.

### 3. Additional *Burst model* Figure

The intention with this figure, is to illustrate that the realized speed of the interferon is connected to the speed of the virus.
The interferon range $r_n > r_v$ typically, Fig. S6 illustrates that the speed gain of the interferon is first realized when $r_v \sim r_n$.

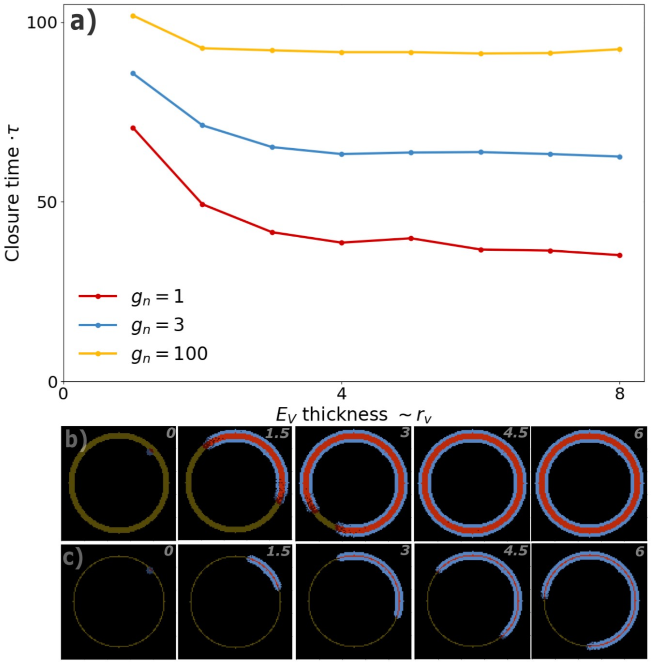


FIG. S6. **Quantifying realized speed of the the interferon in the "Burst-Scenario":** Across all simulations $r_n = 3$ is fixed, and $E_V$ ***state is frozen***, that means $E_V$ **never** goes to $L_V$. This synthetic setup is made to conceptually illustrate that the realized speed of the interferon is partially set by the by the virus, since the interferon is only able to propagate signal within virus exposed tissue.
Panel a) explores the closure time across different interferon strategies. quantifies the closure speed of a circle for different shape parameters $g_n$. The true scaling power of the IFN spreading is first reached when $r_v \sim r_n$. This effect is illustrated that the red curve and the blue curve ($g_n = 1, 3$) saturates at $E_V$-thickness$= r_n = 3$
b), c) shows sample simulations, b) has $E_V$-thickness 3, while c) has $E_V$-thickness 1. Both b),c) has $g_n = 1$. This tells us that the realized speed of an interferon is not only a function of $r_n, g_n$ but also a function on $r_v$. $Speed_N$ increases as $r_v$ approaches $r_n$ from below, and saturates to maximal possible realized speed once $r_v \sim r_n$.
This effect would also be seen in the "*Shedding-Scenario*" if the exposed state where allowed to be stochastic.

### 4. Additional Data Figures

In this section a dynamic intracellular viral RNA time series is shown in fig S7 for the lytic Polio virus in human epithilial cells, and the production rate of the non lytic VSV virus in baby hamster kidny cells.

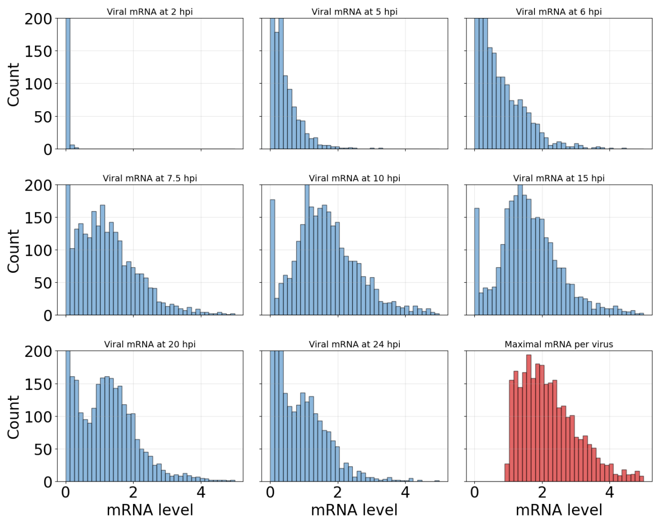


FIG. S7. This shows the concentration of intracellular Polio viral RNA in the human epithilial cells at different time points, using data from [9]. The last distribution shows the maximal intracellular viral RNA at any given time point within the time sample. The experiment is tracked for 24 hours.

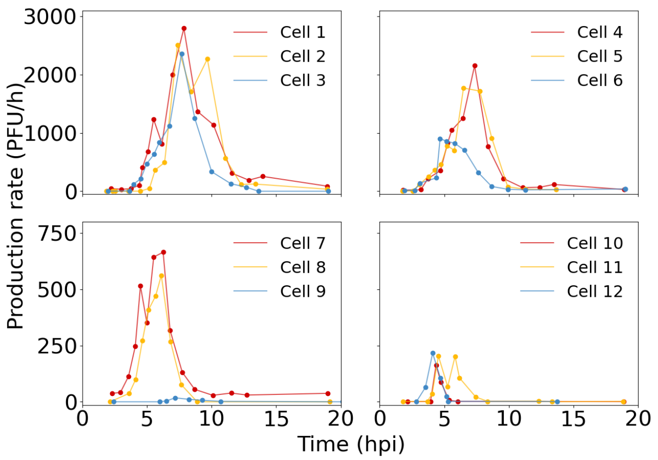


FIG. S8. This is the same data as shown in fig 1 panels b), c), visualized with no fits. The data used is from [16]

.
